# Rare centenarian SIRT6 variants elevate SIRT6 protein levels and resist cellular senescence

**DOI:** 10.1101/2025.09.29.679208

**Authors:** Jiping Yang, HyeRim Han, Xifan Wang, Yizhou Zhu, Haiqi Chen, Yousin Suh

## Abstract

Centenarians provide valuable insights into the biological mechanisms underlying human longevity and potential gerotherapeutic targets. We previously identified two linked missense variants in *SIRT6* that are enriched in Ashkenazi Jewish centenarians. To investigate their functional impact in physiologically relevant cellular contexts, we generated human embryonic stem cells carrying these variants through precise genomic knock-in and differentiated them into somatic lineages. Functional characterization revealed that the centenarian variants endogenously elevated SIRT6 protein levels through weakened interaction with vimentin, and altered SIRT6 enzymatic activities including enhanced mono-ADP-ribosyl transferase activity and reduced deacetylase activity. These variants delayed replicative and progerin-induced cellular senescence, preserving genome stability through maintenance of DNA repair pathways and suppression of transposable element derepression. Moreover, pharmacologically mimicking the centenarian variants using SIRT6 activator Fucoidan-FV partially ameliorated premature aging–associated molecular defects in progeria fibroblasts. Together, our findings demonstrate that rare centenarian variants exert multifaceted effects on SIRT6 and enhance cellular resilience, providing insights for developing geroprotective therapies informed by genetic discoveries in exceptionally long-lived individuals.

## Introduction

Sirtuin 6 (SIRT6) is a nicotinamide adenine dinucleotide (NAD^+^)-dependent enzyme with deacetylase and mono-ADP-ribosyl transferase (mADPr) activities that plays pivotal roles in DNA damage repair, chromatin stability, metabolic regulation, and stress responses^1,2^. SIRT6 overexpression extends mouse lifespan^3,4^ and its double-strand break repair activity correlates with lifespan across species^5^, firmly establishing *SIRT6* as a key longevity gene.

We previously identified two linked missense variants in *SIRT6* (rs201141490-N308K and rs183444295-A313S) that are enriched in a unique centenarian cohort^6^. The resulting centenarian SIRT6 protein (CentSIRT6) exhibits altered mono-ADP-ribosyltransferase and deacetylase activities, and enhances double-strand break repair and cancer cell killing^6^. However, prior protein-centric study has largely relied on ectopic expression of the mutant protein in immortalized cell lines, an approach that may not accurately recapitulate the physiological context in which these variants naturally function. Consequently, the impact of these longevity-associated variants on molecular and cellular outcomes within their endogenous genomic environment remains unclear, underscoring the critical need for physiologically relevant cellular models to better assess their biological roles.

To address this gap, we used CRISPR/Cas9 to introduce the centenarian SIRT6 variants into human embryonic stem cells (hESCs) and differentiated them into somatic cell types for functional characterization. Using this more physiologically relevant cell model, we discovered that centenarian SIRT6 variants elevate SIRT6 protein levels through reduced interaction with vimentin - an effect not observed in ectopic expression systems, and confirmed the previously characterized enzymatic alterations. Functionally, the variants delayed replicative and progerin-induced cellular senescence by preserving genome stability. Notably, treatment with SIRT6 activator Fucoidan-FV, which mimics the centenarian variants by increasing mADPr activity and SIRT6 protein levels, partially rescued premature aging-associated molecular defects in fibroblasts derived from Hutchinson–Gilford progeria syndrome (HGPS) patients. Collectively, these findings illuminate how rare centenarian variants modulate SIRT6 function and cellular resilience, expanding understanding of genetic and molecular basis of human longevity and providing avenues to translate genetic discoveries from long-lived individuals into gerotherapeutic strategies for healthy aging.

## Results

### Centenarian SIRT6 variants elevate SIRT6 protein levels in an endogenous genomic context

To investigate the impact of centenarian SIRT6 variants in an endogenous genome context, we introduced the centenarian variants (rs183444295 and rs201141490) into hESCs (wild-type, WT) using CRISPR/Cas9-mediated gene editing **(Fig. 1A)**. Successfully edited hESCs (centenarian, CENT) maintained pluripotency maker OCT4 expression, comparable to it isogenic controls **(Fig. S1A)**.

**Fig. 1.**
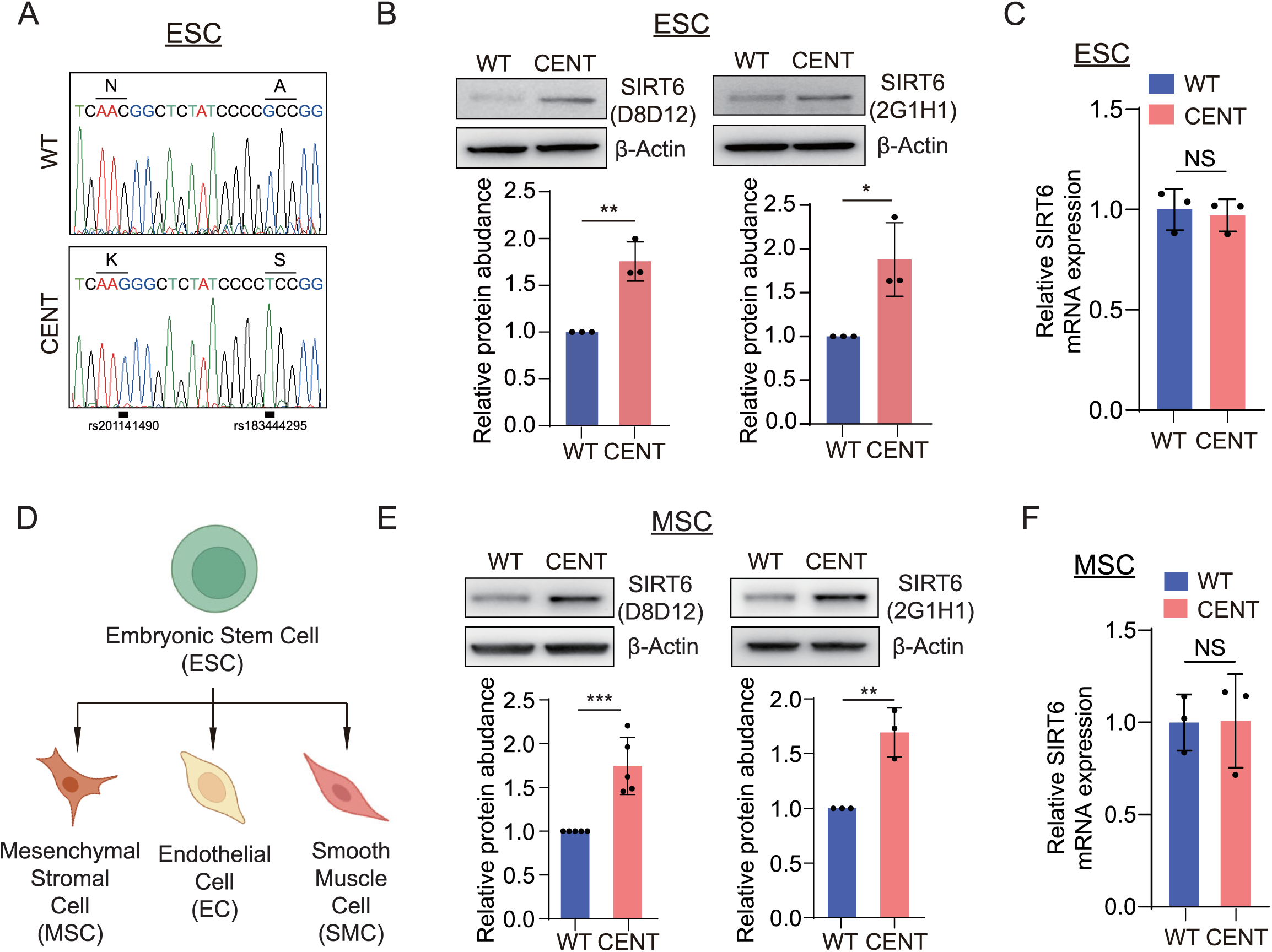
Elevated SIRT6 protein levels in hESCs and hESC-derived hMSCs carrying centenarian SIRT6 variants. **(A)** Sanger sequencing confirming the knock-in of centenarian SIRT6 variants in WT and CENT hESCs. **(B)** SIRT6 protein levels in WT and CENT hESCs measured using two different SIRT6 antibodies (Clone ID: D8D12 and 2G1H1); Protein levels were normalized to β-Actin; data were represented as mean ± SD; n = 3; *, p < 0.05; **, p < 0.01. **(C)** SIRT6 mRNA levels in WT and CENT hESCs, gene expression was normalized to 18s rRNA; data were represented as mean ± SD; n = 3; NS, not significant. **(D)** Schematic showing the hESC-differentiated somatic cell types used in subsequent analyses. **(E)** SIRT6 protein levels in WT and CENT hMSCs; Protein levels were normalized to β-Actin; data were represented as mean ± SD; n = 3; **, p < 0.01; *** p < 0.001. **(F)** SIRT6 mRNA levels in WT and CENT hMSCs, gene expression was normalized to 18s rRNA; data were represented as mean ± SD; n = 3; NS, not significant.

Unexpectedly, CENT hESCs showed significantly elevated SIRT6 protein levels despite unchanged mRNA levels **(Fig. 1B-1C)**. To assess whether this effect was cell-type specific, we differentiated hESCs into human mesenchymal stromal cells (hMSCs), human endothelial cells (hECs) and human smooth muscle cells (hSMCs) **(Fig. 1D)**. Each lineage expressed its respective markers **(Fig. S1B)**, confirming lineage-committed differentiation. Notably, all three CENT-derived lineages showed consistently elevated SIRT6 protein levels as confirmed using two antibodies recognizing distinct SIRT6 epitopes **(Fig. 1E and S1D-E)** but unchanged mRNA levels **(Fig. 1F and S1C)**. Given that hMSCs are well-established cellular model for aging research^7–11^, we focused subsequent functional analyses on this cell type.

### Elevated SIRT6 is not caused by increased nuclear localization

Next, we investigated how the centenarian variants elevate SIRT6 protein levels. Since these centenarian mutations (N308K and A313S) are located proximal to the C-terminal nuclear localization signal (NLS; amino acids 331-353) **(Fig. 2A)**, raising the possibility of altered nuclear import, we examined their effect on subcellular distribution. However, subcellular fractionation revealed elevated SIRT6 in both cytoplasmic and nuclear compartments of CENT hMSCs **(Fig. 2B)**, ruling out enhanced nuclear import.

**Fig. 2.**
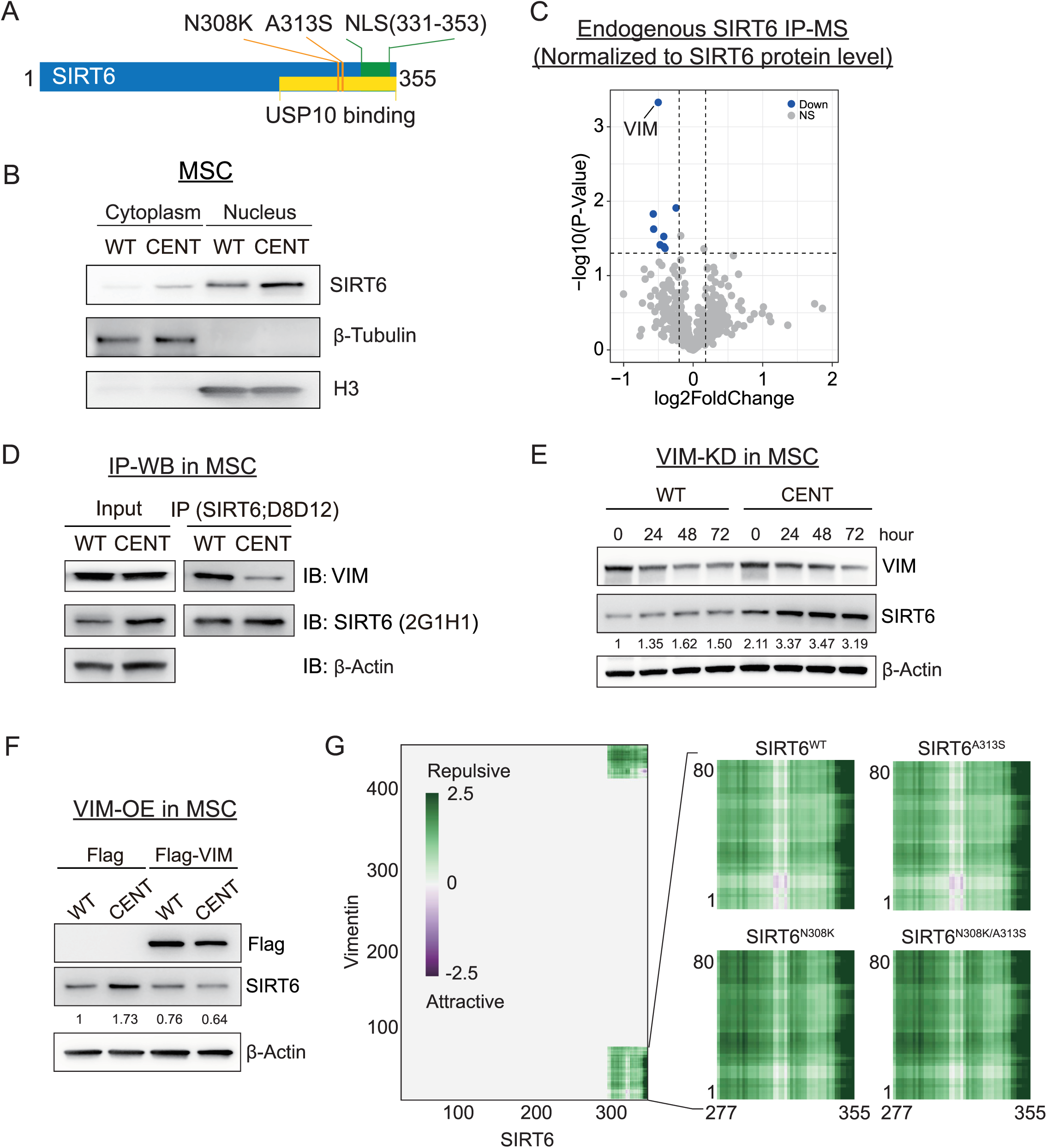
Vimentin is responsible for the elevated SIRT6 protein levels in CENT hMSCs. **(A)** Schematic representation of the known domains in the SIRT6 protein. The green bar denotes the nuclear localization signal (NLS), orange lines indicate the positions of centenarian SIRT6 mutations, and the yellow bar marks the C-terminal region required for USP10 binding. **(B)** SIRT6 protein levels in cytoplasmic and nuclear fractions of WT and CENT hMSCs. **(C)** Volcano plot showing differentially interacting proteins between WT SIRT6 and CentSIRT6 in hMSCs, identified by endogenous SIRT6 IP-MS. Protein abundances were normalized to SIRT6 levels in each pull-down sample. **(D)** Endogenous SIRT6 was immunoprecipitated from WT and CENT hMSC lysates using anti-SIRT6 antibody (D8D12). IP samples and input lysates were analyzed by western blotting with antibodies against SIRT6 (2G1H1) and Vimentin (VIM). Input represents whole-cell lysate prior to immunoprecipitation. **(E)** Protein levels of Vimentin and SIRT6 in WT and CENT hMSCs transfected with siRNAs targeting Vimentin. Protein levels were normalized to β-Actin. **(F)** Protein levels of Flag-VIM and SIRT6 in WT and CENT hMSCs transduced with lentiviruses expressing Flag or Flag-VIM. Protein levels were normalized to β-Actin. **(G)** Predicted interaction between the VIM IDR and the C-terminal IDR of SIRT6 using FINCHES. The right panel showing the effects of different SIRT6 mutations on these predicted interactions.

### Elevated SIRT6 is independent of its stabilizer USP10

SIRT6 stability is regulated by the ubiquitin-proteasome system through interactions with E3 ubiquitin ligases^12,13^ and deubiquitinating enzymes (DUBs)^14^. Among its established interactors, the DUB USP10 stabilizes SIRT6 by binding its C-terminal region^14^ - the region harboring the centenarian mutations **(Fig. 2A)**. To test whether the elevated SIRT6 is mediated by USP10, we performed USP10 knockdown using siRNAs in both WT and CENT hMSCs. While USP10 knockdown reduced SIRT6 protein levels in WT hMSCs, SIRT6 protein levels in CENT hMSCs remained unchanged **(Fig. S2A)**, indicating that the increased SIRT6 levels in CENT hMSCs is independent of USP10.

### Vimentin mediates the increased SIRT6

To unbiasedly identify mediators contributing to elevated SIRT6, we performed endogenous SIRT6 immunoprecipitation (IP) followed by tandem mass tag (TMT) - labeled mass spectrometry (MS) in WT and CENT hMSCs. Strikingly, the most significantly altered SIRT6 interactor was vimentin (VIM), an intermediate filament protein **(Fig. 2C)**. Endogenous centSIRT6 protein exhibited weaker interaction with vimentin compared to WT-SIRT6 **(Fig. 2C**–**2D)**. Moreover, vimentin knockdown by siRNAs increased SIRT6 protein levels in both WT and CENT hMSCs **(Fig. 2E)**, whereas vimentin overexpression reduced SIRT6 levels in CENT hMSCs to those levels in WT hMSCs **(Fig. 2F)**. These results demonstrate that vimentin mediates the increased SIRT6 in CENT hMSCs.

To gain insight into how the centenarian mutations alter the SIRT6–vimentin interaction, we focused on their intrinsically disordered regions (IDRs) ^2,15^, which are known to mediate protein–protein interactions^16^. AlphaFold^17^ predictions revealed that SIRT6 harbors a C-terminal IDR (amino acids 298-355) and vimentin contains two IDRs at N-terminal (amino acids 1-89) and C-terminal (amino acids 409-422) regions **(Fig. S2B)**. To assess whether the IDRs of SIRT6 and vimentin have binding potential, we used FINCHES^18^, a sequence-based tool that can predict chemical-specific intermolecular interactions driven by disordered regions. We found SIRT6’s C-terminal IDR encompassing the centenarian mutations showed potential to bind both the N-terminal and C-terminal IDRs of vimentin **(Fig. 2G)**. Introduction of the two centenarian mutations reduced the attraction between N-terminal IDRs of vimentin and SIRT6, consistent with our experimental findings **(Fig. 2G)**. Notably, only the N308K mutation, not A313S which contributes more to enhanced mADPr activity^6^, weakened the predicted SIRT6-vimentin interaction **(Fig. 2G)**, suggesting N308K is responsible for the reduced SIRT6-vimentin interaction and the consequent increase in SIRT6 protein levels.

### Centenarian SIRT6 variants alter enzymatic activities in a cellular context

Given that CentSIRT6 protein displays weaker deacetylase activity but stronger mADPr activity in biochemical assays^6^, we next assessed these activities in a cellular context using isogenic hMSCs. Despite having higher SIRT6 protein, CENT hMSCs showed increased histone acetylation at H3K9, H3K18, and H3K27 compared to WT hMSCs **(Fig. S3A)**, confirming the reduced deacetylase activity of CentSIRT6. Additionally, we observed an increased proportion of mADP-ribosylated SIRT6 protein in CENT hMSCs **(Fig. S3B)**, indicating enhanced mADPr activity of CentSIRT6. These results reveal that centenarian SIRT6 variants alter enzymatic activities in a cellular context, consistent with previous biochemical findings.

### Centenarian SIRT6 variants moderately alter the transcriptome of young hMSCs

To investigate the molecular changes caused by centenarian SIRT6 variants, we performed transcriptome analysis on WT and CENT hMSCs at early passage. Differentially expressed gene (DEG) analysis demonstrated that replacement of two nucleotides altered the MSC transcriptome, revealing 52 down-regulated and 72 up-regulated DEGs (|Fold Change| > 2, adjusted p value < 0.05) in CENT versus WT hMSCs **(Fig. 3A)**. Gene ontology analysis showed that these DEGs were significantly enriched in extracellular matrix (ECM) terms **(Fig. 3B)**.

**Fig. 3.**
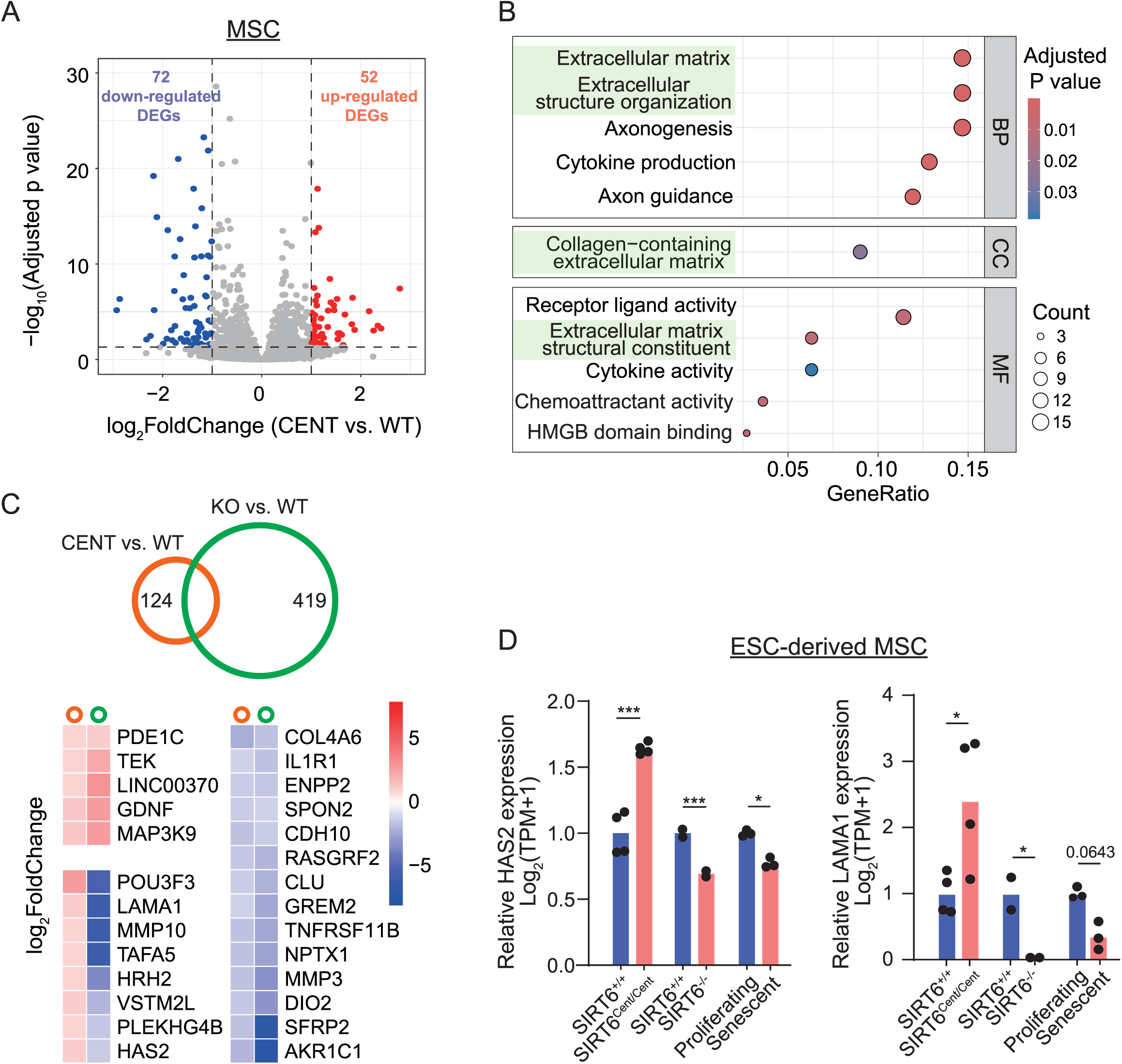
Moderate transcriptomic changes between WT and CENT hMSCs at early passage. **(A)** Volcano plot showing the differentially expressed genes (DEGs) identified by RNA-seq of WT and CENT hMSCs at early passage (P3). **(B)** Gene ontology (GO) analysis of DEGs between WT and CENT hMSCs. Extracellular matrix (ECM)-associated GO terms were highlighted in green. **(C)** Venn diagram showing the overlap of DEGs identified in CENT vs. WT hMSCs and SIRT6-KO vs. WT hMSCs. The lower panel presents the overlapping genes along with their expression changes in both comparisons. **(D)** RNA-seq expression profiles of ECM-related genes *HAS2* and *LAMA1* in CENT vs. WT hMSCs, SIRT6-KO vs. WT hMSCs, and proliferating vs. senescent hMSCs. *, p < 0.05; ***, p < 0.001. Normalized expression values (TPM: Transcripts Per Million) were used for plotting; data were represented as mean.

To identify genes potentially regulated directly by SIRT6, we integrated transcriptome data from SIRT6-knockout hMSCs^19^, which revealed 419 DEGs relative to SIRT6-WT hMSCs **(Fig. 3C)**. Among these, 27 genes overlapped with those affected by the centenarian variants, showing both concordant and opposite expression changes **(Fig. 3C)**, suggesting that the effects of centenarian SIRT6 variants are more complex than simple up- or down-regulation of SIRT6. Notably, the ECM genes *HAS2* and *LAMA1*, which were downregulated in SIRT6-knockout hMSCs and during MSC senescence, were significantly upregulated in CENT hMSCs **(Fig. 3D)**. Interestingly, *HAS2*, encoding hyaluronan synthase 2, is an ECM component that has been linked to improved healthspan^20^.

We also examined genes in pathways previously linked to SIRT6 including glycolysis^21,22^, lipogenesis^22^, autophagy^23,24^, oxidative stress^19^ and cancer^2^, but most showed no significant differences between WT and CENT hMSCs **(Fig. S4)**. Taken together, the SIRT6 variants exerted only moderate effects on the transcriptome of young hMSCs, which remained largely homeostatic and minimally stressed.

### Centenarian SIRT6 variants delay replicative senescence

Given that SIRT6 is highly dynamic and stress-responsive^25^, we next investigated the centenarian variants in hMSCs under stress, examining their effects on cellular aging through replicative senescence induced by serial passaging **(Fig. S5A)**. Early-passage WT and CENT hMSCs showed similar proliferation (Ki67+ cells) and senescence-associated β-galactosidase (SA-β-gal) activity **(Fig. S5B-S5C)**. However, at late passages, CENT hMSCs exhibited a higher proportion of Ki67-positive cells and fewer SA-β-gal-positive cells **(Fig. S5B-S5C)**, demonstrating delayed senescence compared to WT hMSCs.

### Centenarian SIRT6 variants confer resistance to progerin-induced stress

Since SIRT6 maintains chromatin organization, DNA repair-cellular processes disrupted by progerin^26^, we next evaluated the impact of the centenarian SIRT6 variants in a more physiologically relevant context: Hutchinson-Gilford Progeria Syndrome (HGPS), an premature aging disorder. HGPS is caused by progerin, a toxic isoform of *LMNA* caused by a splicing mutation. We generated WT and CENT hMSCs stably expressing GFP-progerin (Progerin-hMSCs)^27^. Both lines showed comparable progerin levels **(Fig. 4A)**, with CENT Progerin-hMSCs retaining elevated SIRT6 protein levels **(Fig. 4B)**. Colony formation assays revealed enhanced proliferative potential of CENT Progerin-hMSCs **(Fig. 4C)**. Moreover, centenarian variants strongly ameliorated multiple cellular aging phenotypes **(Fig. 4D)**, as evidenced by a higher percentage of Ki67-positive cells **(Fig. 4E)**, reduced SA-β-gal activity **(Fig. 4F)**, decreased expression of aging markers P16, P21, senescence-associated secretory factor (SASP) gene IL6, and increased levels of Lamin B1 **(Fig. 4G)**.

**Fig. 4.**
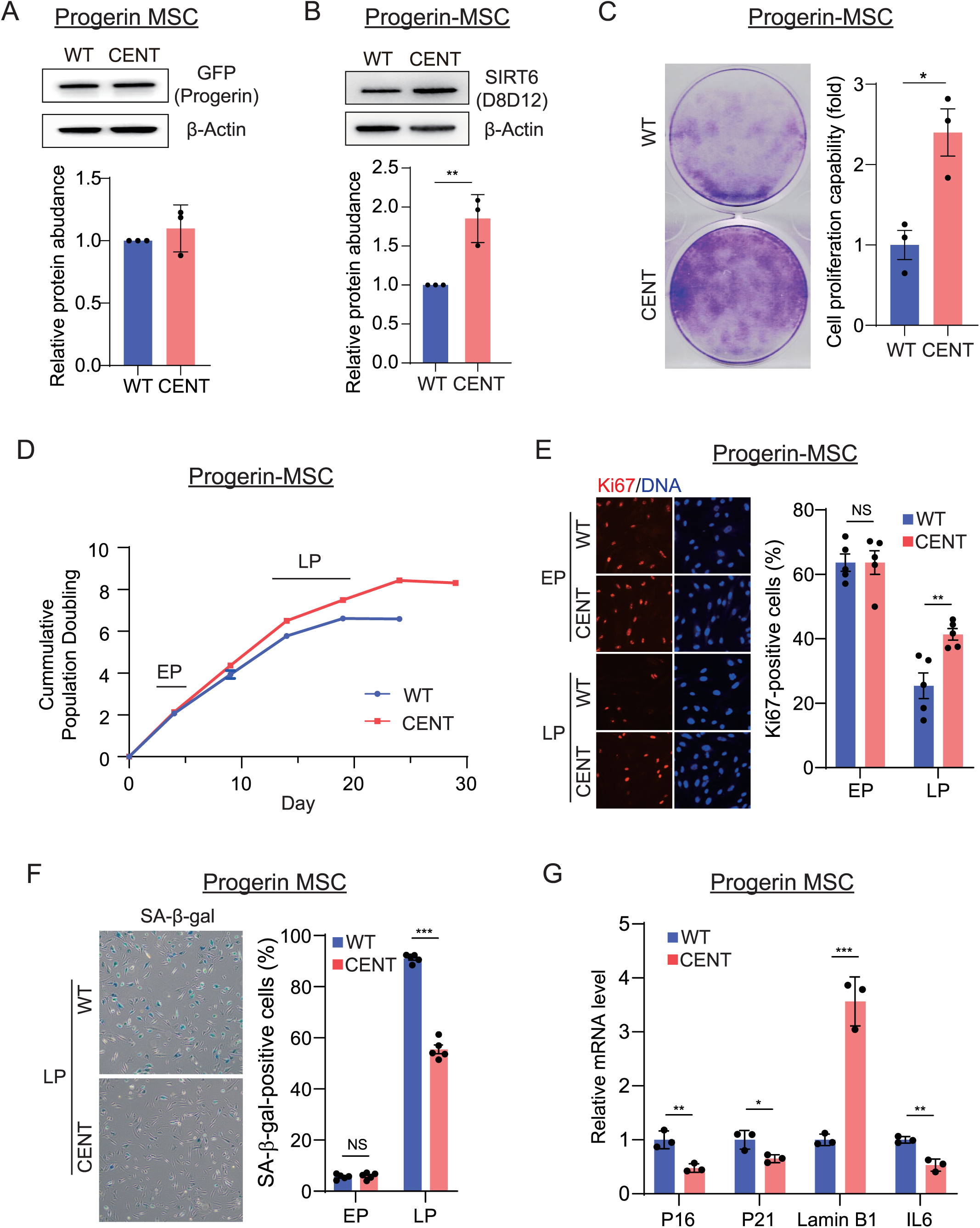
Centenarian SIRT6 variants confer resistance to progerin-induced stress. **(A)** Progerin protein levels in WT and CENT hMSCs transduced with GFP-progerin lentiviruses (Progerin-MSCs). Protein levels were normalized to β-Actin; data were represented as mean ± SD; n = 3; NS, not significant. **(B)** SIRT6 protein levels in WT and CENT Progerin-MSCs. Protein levels were normalized to β-Actin; **, p < 0.01. **(C)** Clonal formation analysis of WT and CENT Progerin-MSCs; data were represented as mean ± SD; n = 3. *, p < 0.05. **(D)** Growth curves of WT and CENT Progerin-MSCs. EP (early passage, 2 passages post-Progerin overexpression and sorting); LP (late passage, 4-5 passages post-Progerin overexpression and sorting), data were represented as mean ± SD, n = 2. **(E)** Immunofluorescence (IF) staining of Ki67 in WT and CENT Progerin-MSCs at EP and LP. Quantification based on 5 independent images capturing over 500 nuclei; data were represented as mean ± SD; NS, not significant; **, p < 0.01. **(F)** SA-β-gal staining in WT and CENT Progerin-MSCs at EP and LP. Quantification based on 5 independent images capturing over 500 nuclei; data were represented as mean ± SD; NS, not significant; ***, p < 0.001. **(G)** Relative mRNA levels of P16, P21, Lamin B1 and IL6 in WT and CENT Progerin-MSCs at LP; gene expression was normalized to 18s rRNA; data were represented as mean ± SD; n = 3, *, p < 0.05; **, p < 0.01; ***, p < 0.001.

### SIRT6 centenarian variants maintain genome stability under progerin-induced stress

Next, we investigated the molecular mechanisms by which the centenarian SIRT6 variants confer protection against progerin. Transcriptome analysis of WT and CENT Progerin-hMSCs identified 312 down-regulated and 143 up-regulated DEGs (|Fold Change| > 2, adjusted p value < 0.05) in CENT versus WT hMSCs **(Fig. 5A)**. Compared to early passage cells, the increased number of DEGs under progerin-induced stress demonstrated that SIRT6 is stress-responsive. Gene set enrichment analysis revealed significant alterations in pathways related to cell cycle regulation and genome stability including not only DNA damage repair such as double-strand break (DSB) and base excision repair (BER) but also higher-order structures such as chromosome and telomere maintenance **(Fig. 5B**–**5C)**. At early passage without progerin, many DNA repair genes showed comparable expression in WT and CENT hMSCs. With progerin expression, the expression of these genes was markedly declined in WT hMSCs but largely preserved in CENT MSCs **(Fig. 5D)**, suggesting that the centenarian SIRT6 variants primarily ameliorate progerin-induced the stress through enhancing genome stability.

**Fig. 5.**
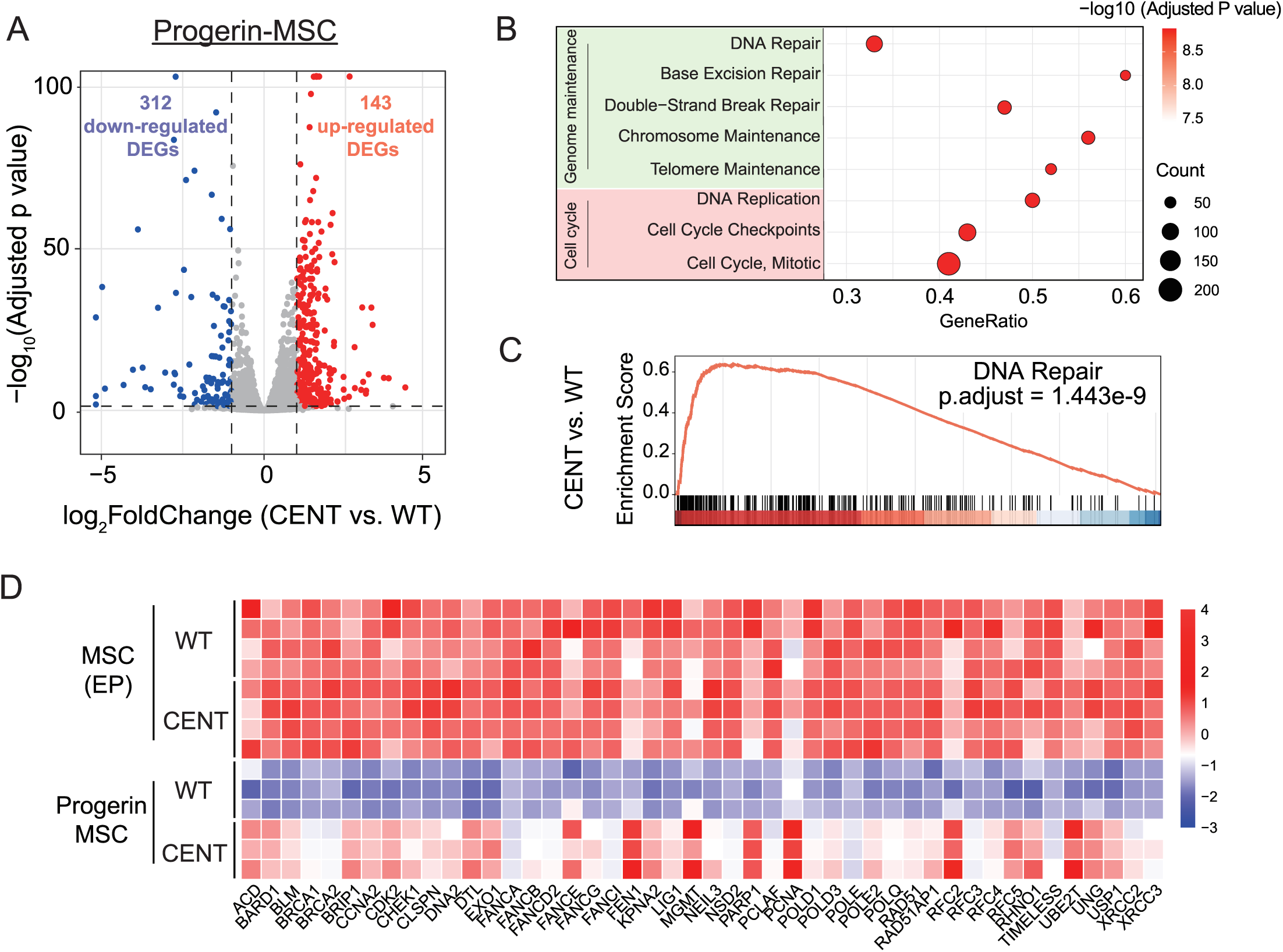
SIRT6 centenarian variants maintain genome stability under progerin-induced stress. **(A)** Volcano plot showing the differentially expressed genes (DEGs) identified by RNA-seq of WT and CENT Progerin-MSCs at 3 passages post-Progerin overexpression and sorting. **(B)** Gene set enrichment analysis (GSEA) revealing pathways altered by centenarian SIRT6 variants were enriched in cell cycle regulation and genome maintenance. **(C)** GSEA enrichment plot showing downregulation of genes involved in DNA repair pathways in CENT Progerin-MSCs compared to WT Progerin-MSCs. **(D)** Heatmap showing the expression of DNA repair genes in hMSCs with or without Progerin-induced stress.

### Centenarian SIRT6 variants suppress progerin-Induced TE derepression

Progerin triggers heterochromatin loss^28^, leading to the derepression of transposable elements (TEs) such as LINE1^29^ and HERVs^30^. SIRT6 maintains TE silencing, but this fails in senescent cells or aged tissues, allowing TEs to become transcriptionally active and contribute to genomic instability^31,32^. We therefore examined whether centenarian SIRT6 variants could mitigate progerin-induced TE reactivation. We found minimal changes in early passage **(Fig. 6A)**, but upon progerin-induced stress, CENT hMSCs showed better maintained suppression **(Fig. 6A)**, particularly of LINE1 elements **(Fig. 6B)**, compared to WT hMSCs. We further confirmed reduced LINE1 ORF1p and ORF2p proteins **(Fig. 6C)**, both translated from LINE1 RNAs in Progerin-MSCs carrying centenarian SIRT6 variants. Additionally, the majority of non-LINE TEs also exhibited reduced expression in CENT hMSCs compared to WT hMSCs following progerin overexpression **(Fig. 6D)**, suggesting globally preserved genome integrity conferred by centenarian SIRT6 variants.

**Fig. 6.**
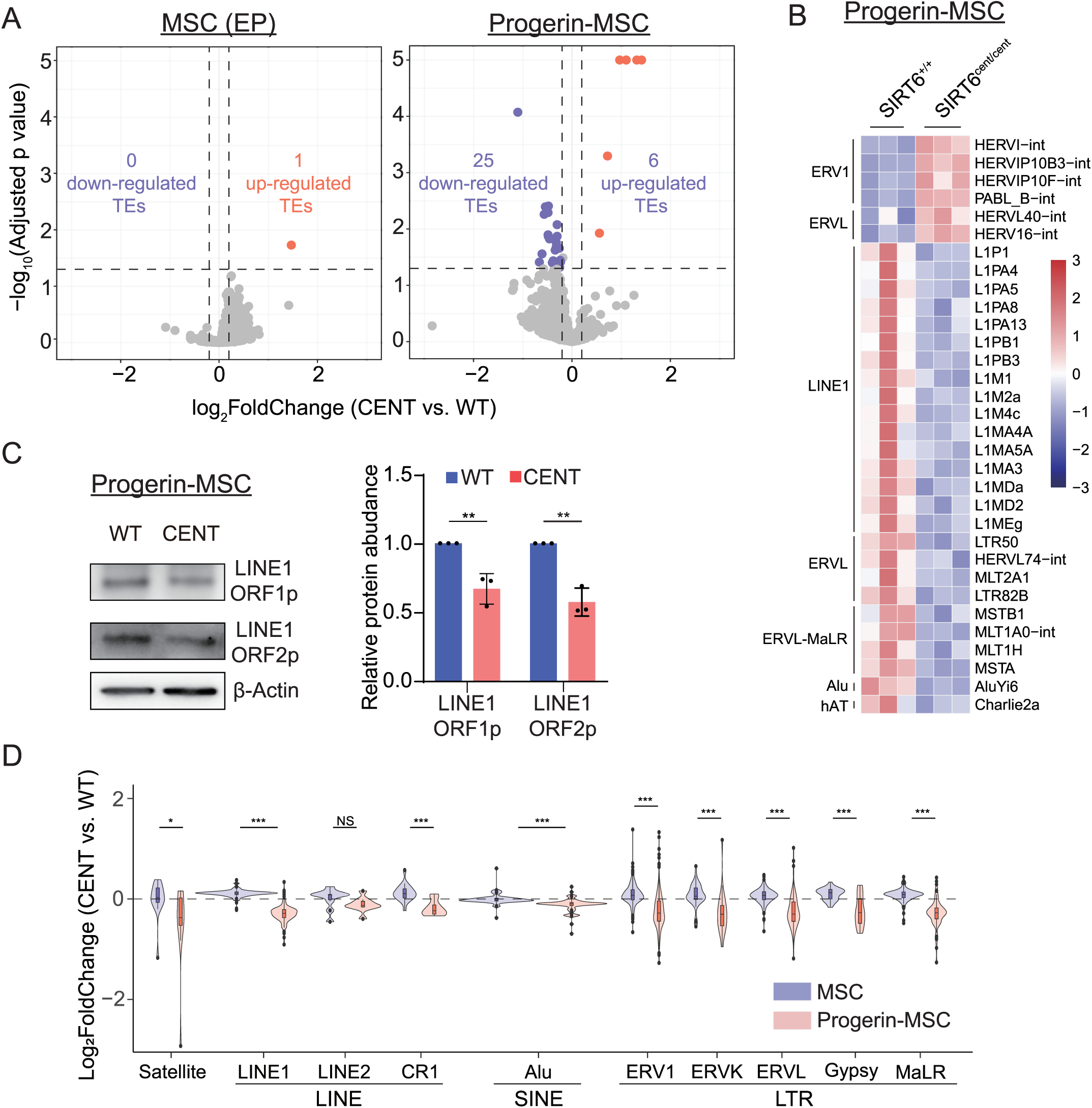
Centenarian SIRT6 variants suppress progerin-Induced TE derepression. **(A)** Volcano plot showing the differentially expressed transposable elements (DETEs) identified in EP hMSCs (Left) and Progerin-MSCs (Right). **(B)** Heatmap displaying the expression levels of DETEs in Progerin-MSCs. **(C)** Protein levels of LINE1-ORF1p and LINE1-ORF2 in WT and CENT Progerin-MSCs. Protein levels were normalized to β-Actin; data were represented as mean ± SD; n = 3; **, p < 0.01. **(D)** Violin plot showing the fold changes of categorized TEs in EP MSC and Progerin-MSCs. NS, not significant; *, p < 0.05; **, p < 0.01; ***, p < 0.001.

### SIRT6 activator Fucoidan-FV represses LINE1 in progeria patient cells

Fucoidan from *Fucus vesiculosus* (Fucoidan-FV) was recently identified as a SIRT6 activator that elevates SIRT6 levels and enhances mADPr activity^33,34^, mimicking the effects of centenarian SIRT6 variants. To investigate its potential in alleviating progeria, we performed a dose-dependent screening in HGPS-patient derived fibroblasts using LINE1 ORF1p expression as a readout **(Fig. 7A)**. Treatment with 50 and 100 μg/mL Fucoidan-FV reduced LINE1 ORF1p protein levels compared to no treatment **(Fig. 7A**–**7B)**. Fucoidan-FV had no detectable effect on endogenous progerin, other senescence-associated markers including P16, P21, Lamin B1, or the heterochromatin marker H3K9me3 **(Fig. 7B)**. Fucoidan-FV also did not alter the proportion of SA-β-gal–positive or Ki67-positive cells **(Fig. 7C**–**7D)**. These results suggest that Fucoidan-FV can selectively mitigate senescence-associated molecular defects, particularly LINE1 reactivation, although its impact on broader senescence phenotypes is limited, likely due to the near-senescent state of the patient-derived cells as evidenced by < 5% Ki67 positivity.

**Fig. 7.**
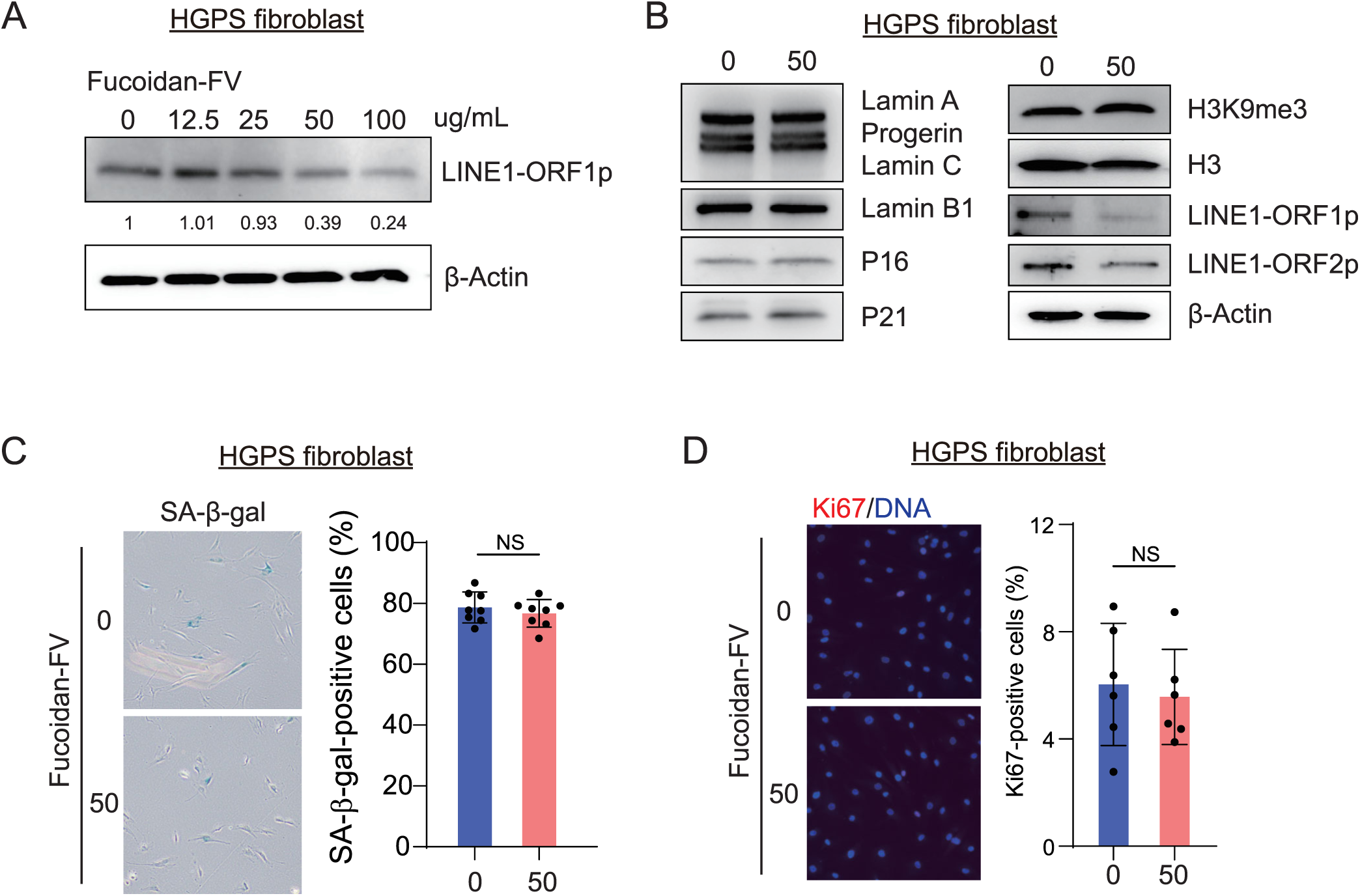
Treatment with SIRT6 activator Fucoidan-FV downregulates LINE1 in progeria patient cells. **(A)** LINE1-ORF1p protein levels in human progeria fibroblasts after the treatment with Fucoidan-FV (0-100 μg/mL). Protein levels were normalized to β-Actin. **(B)** Protein levels of senescence markers in HGPS patient fibroblasts treated with or without 50 μg/mL Fucoidan-FV. H3 and β-Actin are used as loading controls. **(C)** SA-β-gal staining in HGPS patient fibroblasts treated with or without 50 μg/mL Fucoidan-FV. Quantification based on 8 independent images capturing over 200 nuclei; data were represented as mean ± SD; NS, not significant. **(D)** IF staining of Ki67 in HGPS patient fibroblasts treated with or without 50 μg/mL Fucoidan-FV. Quantification based on 6 independent images capturing over 200 nuclei; data were represented as mean ± SD; NS, not significant.

## Discussion

Centenarians, who live not only longer but also healthier, hold the key to access longevity and healthy aging. Genetic association studies have identified variants robustly associated with human longevity, thereby advancing our understanding of human aging and longevity^35^. However, the translation of these findings from long-lived individuals remains largely unexplored. A major barrier is the lack of variant-to-function studies elucidating their functional roles^36^. In this study, we applied variant knock-in in human embryonic stem cells followed by directed differentiation, representing a reliable and physiologically relevant *in vitro* method to assess the function of rare centenarian missense variants.

This approach demonstrated advantages in two key aspects. First, unlike overexpression systems, the knock-in strategy maintains the gene’s natural regulatory environment, ensuring physiological transcription, splicing, and protein expression levels. By introducing the variant precisely into the endogenous locus, knock-in models enable direct comparison with isogenic wild-type controls, providing clear causal evidence for how the variant impacts gene functions and cellular outcomes. Using this approach, we uncovered a novel mechanism whereby centenarian SIRT6 variants increase SIRT6 protein levels through weakened vimentin interaction-effects that could not be detected through conventional ectopic expression, alongside the previously characterized enzymatic alterations. Second, unlike immortalized cell lines with infinite *in vitro* lifespan, hESC-derived somatic cells provide a more physiologically relevant system for evaluating variant impacts on cellular senescence and offer potential for exploring cell-type specific effects. hESC-derived hMSCs represent normal diploid cells and are widely used as a cellular aging model to evaluate the effects of genes^11^ or compounds^37^ on senescence. Our study demonstrated that centenarian SIRT6 variants delay cellular aging primarily by enhancing genome stability.

The finding that vimentin regulates SIRT6 stability through IDR-mediated interactions reveals a novel proteostasis mechanism. Vimentin, traditionally recognized as an intermediate filament protein providing structural support, has more recently been implicated in autophagy^38^ and vesicle trafficking^39,40^, underscoring its role in proteostasis. Notably, vimentin has been reported to undergo liquid–liquid phase separation (LLPS) to form droplets^15^, a process that depends on its IDR. The precise mechanism by which Vimentin stabilizes centSIRT6 remains elusive, but likely involves LLPS-dependent vesicle trafficking, with SIRT6 serving as a cargo.

Our study highlights three major translational opportunities to improve human healthspan. First, given that SIRT6 overexpression extends healthspan and lifespan in mice, systematic delivery of CentSIRT6 via safe delivery approaches, such as Adeno-associated virus (AAV), could represent a promising gene therapy to combat aging. Second, centenarian variants can be leveraged to genetically enhance stem cell function for cell therapy, a concept similar to recent studies introducing artificial mutations in longevity gene *FOXO3* to generate senescence-resistant MSCs (SRCs) which counter aging in primates^41^. Compared to artificial mutations where the site also has mutation in cancer cells, centenarian variants carry minimal safety concerns as individuals naturally harbor these variants for a century and maintain longer and healthier lives. Third, our study can guide the development of SIRT6 activators for geroprotection. Existing SIRT6 activators primarily enhance deacetylase activity^2^, but our data suggest boosting mADPr activity may be more critical for extending human healthspan and lifespan.

The centenarian SIRT6 variants confer multifaceted effects on SIRT6 protein. This complex gain-of-function illustrates why rare missense variants cannot be phenocopied by conventional gene overexpression or knockdown strategies commonly used for studying GWAS variants. Our variant knock-in and hESC differentiation paradigm partially addresses the variant-to-function challenges, primarily revealing molecular and cellular changes such as stress response and senescence. Nevertheless, gaps remain in understanding tissue-specific consequences of the variants and their overall impact on healthspan and lifespan. Future studies should leverage *ex vivo* human hESC-derived organoids to directly interrogate variant effects in physiologically relevant tissue contexts. Additionally, given that these variants are not conserved in rodents, humanized SIRT6 mice or non-human primates represent optimal models to evaluate their impact on healthspan and lifespan.

In conclusion, this study provides a framework for functionally characterizing rare centenarian variants and translating genetic discoveries into therapeutic interventions. This functional genomics study on centenarian SIRT6 variants demonstrate how natural occurring human mutations can guide the development of strategies to enhance healthspan and combat age-related decline.

## Methods and Materials

### Antibodies

Antibodies for western blotting (WB) and immunofluorescence (IF) staining were obtained from the following sources.

BD Bioscience: CD73-PE (561014), CD90-FITC (555595), CD105-APC (562408), CD31-FITC (555445), P16 (550834);

Santa Cruz Biotechnology: OCT4 (sc-5279), β-Actin (sc-47778), SIRT6 (2G1H1, sc-517196), GFP (sc-9996), β-Tubulin (sc-5274), VIM (sc-6260), Lamin A/C (sc-376248);

Millipore Sigma: H3K9Ac (07-352), SMA (A5228), Flag (F1804), LINE1-ORF1p (MABC1152);

Cell Signaling Technology: SIRT6 (D8D12, 12486), USP10 (5553), H3 (9715), P21 (2947); Abcam: H3K18Ac (ab1191), H3K9me3 (ab8898), Lamin B1 (ab133741), LINE1-ORF2p (ab106004);

Active Motif: H3K27Ac (39034);

Thermo Fisher: Ki67-APC (17-5699-42);

Bio-Rad: Mono-ADP-Ribose (AbD33205).

For endogenous SIRT6 Immunoprecipitation (IP), SIRT6 (D8D12, 12486) antibodies were used in IP, SIRT6 (2G1H1, sc-517196), VIM (sc-6260) and Mono-ADP-Ribose (Bio-Rad, AbD33205) antibodies were used in immunoblotting (IB).

### Cell culture

H7 human embryonic stem cells (ESC) (WA07, WiCell) were maintained on Matrigel (BD Biosciences) in mTeSR Plus medium (STEMCELL Technology).

MSCs were culture in αMEM (Gibco) with 10% fetal bovine serum (FBS, Hyclone or GeminiBio), and 1 ng/mL bFGF (Thermo Fisher).

Endothelial cells were cultured in EGM-2 endothelial cell growth medium (Lonza CC-3162) supplemented with 10nM SB431542 (Selleck), 50 ng/mL VEGF (Peprotech) and 20ng/mL bFgF (Thermo Fisher).

Smooth muscle cells were cultured in N2B27 Medium supplemented with 2ng/mL Activin A (Thermo Fisher) and 2ug/mL Heparin (STEMCELL Technology).

Fibroblasts isolated from Hutchinson–Gilford Progeria Syndrome (HGPS) patients (Coriell, AG11513) were cultured in DMEM (Gibco) with 15% fetal bovine serum (FBS, Hyclone). Fucoidan derived from *Fucus vesiculosus* was purchased from Sigma-Aldrich (F8190) and kindly provided by Paul D. Robbins (University of Minnesota).

All cell culture media were supplemented with 1% penicillin/streptomycin (Gibco) and 50ug/mL Normocin (InvivoGen) to prevent cells from microbial contaminations. Cultured cells are routinely tested for mycoplasma contamination using Mycoplasma PCR Detection Kit (ABM).

### Variant editing at *SIRT6* locus

CRISPR/Cas9-mediated knock-in was performed using Alt-R CRISPR-Cas9 System (IDT) and HDR Donor Oligos. Cas9 nuclease, SIRT6 gRNA (GAATCTCCCACCCGGATCAA) targeting double variant site and single-strand DNA oligo donors (ssODNs) carrying centenarian SIRT6 variants were ordered from IDT. To generate centenarian SIRT6 variants knock-in hESCs, 2×10^5 individualized hESCs were resuspended in 10 μL Resuspension Buffer R (Invitrogen) containing CRISPR ribonucleoproteins (Cas9 protein+gRNA) and ssODNs and were then electroporated using NEON Transfection System (Invitrogen). After electroporation, cells were seeded on Matrigel-coated plates in mTeSR plus with 1x RevitaCell Supplement (Gibco). After 48h expansion, cells were dissociated by Accutase and 10,000 cells were seeded on CytoSort™ Array (10,000 microwells, CELL Microsystem). Once cells were attached, microwells containing single colony were automatically picked and transferred to 96-well plate by CellRaft AIR System (CELL Microsystem). The expanded clones on 96-well plate were genotyped by TaqMan genotyping assay (rs201141490, rs183444295, Thermo Fisher) and further confirmed by Sanger sequencing of PCR products amplified using primers AGCCTCACCTCTGGACAACACAGCAA and TCGTCAACCTGCAGCCCACCAAGCAC.

### Mesenchymal stromal cell differentiation

hMSCs were differentiated from hESCs as previously described with minor modification^42^. Briefly, embryoid bodies were first produced using AggreWell (STEMCELL Technology) and were then plated on Matrigel-coated plates in hMSC differentiation medium (αMEM (Gibco), 10% fetal bovine serum (GeminiBio), 1% penicillin/streptomycin (Gibco), 10 ng/mL bFGF (Thermo Fisher) and 5 ng/mL TGFβ (Thermo Fisher)) for around 10 days till fibroblast-like cells were confluent. These fibroblast-like cells were maintained in hMSC culture medium on Gelatin-coated 10cm dishes for two passages and were further sorted by the BD FACSAria II cell sorter to purify CD73/CD90/CD105 tri-positive hMSCs.

### Endothelial cell differentiation

Differentiation of hESCs into hECs was performed as previously described with minor modifications^43^. Briefly, hESCs were cultured in mTeSR1-Plus media (STEMCELL Technology) for one day and then in M1 medium, containing IWP2 (3 mM, Selleck), BMP4 (25 ng/ml, Peprotech), CHIR99021 (3 mM, Selleck) and bFGF (4 ng/ml, Thermo Fisher), for three days. The following day, M1 medium was removed and replaced with M2 medium with the addition of VEGF (50 ng/ml, Peprotech), bFGF (20 ng/ml, Thermo Fisher) and IL6 (10 ng/ml, Peprotech) to promote endothelial cell emergence for another three days. The differentiated adherent cells were harvested using TrypLE (GIBCO), labeled with CD144 (VE-Cadherin) MicroBeads (130-097-857, Miltenyi Biotec), and separated by OctoMACS™ Separator (Miltenyi Biotec).

### Growth curve assay

Cell population doubling was determined as previously described^42^. Briefly, hMSCs were serially passaged and the number of cells was counted. Population doubling per passage was calculated as log2 (number of cells harvested/number of cells seeded). Cumulative population doublings of the cells were calculated and plotted against culture days.

### Senescence-associated β-galactosidase (SA-β-gal) staining

The SA-β-gal staining of hMSCs was conducted as previously described^42^. Briefly, cells were washed with PBS and fixed at room temperature for 5 minutes using a fixation buffer containing 2% formaldehyde and 0.2% glutaraldehyde. After fixation, cells were washed with PBS and incubated overnight at 37□°C with freshly prepared staining solution (1 mg/mL X-gal, 40 mM citric acid/sodium phosphate pH 6.0, 5 mM potassium ferrocyanide, 5 mM potassium ferricyanide, 150 mM NaCl, 2 mM MgCl_2_). Images were acquired using the EVOS Cell Imaging System (Thermo Fisher). SA-β-gal-positive and total cell numbers were quantified using ImageJ, and the percentage of SA-β-gal-positive cells was calculated for statistical analysis.

### Immunoprecipitation

For endogenous co-immunoprecipitation assays, WT and CENT hMSCs were lysed in ice-cold IP buffer composed of 50 mM Tris-HCl (pH 7.5), 150 mM NaCl, 1 mM EDTA, 1% NP-40, and 5 mM EDTA, supplemented with 1× Protease and Phosphatase Inhibitor Cocktail (Roche). Lysates were cleared by centrifugation at 13,000 rpm for 15 minutes at 4□°C and further quantified using BCA Kit (Thermo Fisher). Equal amounts of lysates from WT and CENT hMSCs were pre-cleared with Pierce Protein A/G Plus Agarose (Thermo Fisher) for 2 hours at 4□°C with rotation. Agarose were separated by centrifugation at 13,000 rpm for 5 minutes at 4□°C. The supernatant was transferred to fresh tubes and incubated with the SIRT6 antibodies (D8D12, Cell Signaling Technology) overnight at 4□°C under constant rotation. The next day, Pierce Protein A/G Plus Agarose (Thermo Fisher) were added to bind antibodies for 2 hours. The beads were washed five times with ice-cold IP buffer and interacting proteins were eluted using Elution Buffer (100 mM Tris-HCl, pH 7.5, 1% SDS) using thermomixer at 65□°C for 10 minutes. Eluted samples were analyzed by western blotting or sent to Innomics for mass spec analysis.

### Mass spectrometry

TMT quantitation on SIRT6-IP samples was performed by Innomics. Samples were first assayed for protein content using the BCA method and then processed using the S-Trap MS sample prep device (PROTIFI) according to the manufacturer’s protocol. Briefly, samples were first reduced with dithiothreitol (DTT) and alkylated with iodoacetamide (IAM) and the resulting samples were digested in the Strap with Trypsin/Lys-C overnight. Digested samples were eluted and then dried by Speed-Vac. The dried samples were reconstituted with 100mM TEAB pH 8.5. All samples were added with their assigned TMT channels. The TMT-added samples were incubated at room temperature for 1 hour for labelling. About 1% of each TMT-labelled sample was aliquoted and mixed with 1% formic acid. TMT-labelled composite was loaded onto the nano LC-MS/MS system for Label Check. After the Label Check was passed, all samples were dried by Speed-Vac and reconstituted with 2% formic acid. The samples were combined into one mixture. The resulting mixture was further desalted using EVOLUTE® EXPRESS ABN according to the manufacturer’s protocol (Biotage). The eluted samples were dried by SpeedVac, reconstituted with high-pH reversed-phase buffer A, and fractioned via offline HPLC fraction. Fraction samples were reconstituted with mobile phase A and loaded onto the nano LC-MS/MS system. About 1uL of each reconstituted sample was injected for analysis with TMT method. TMT quantification was performed with Proteome Discoverer 2.5 (Thermo Fisher).

### Western blotting

Cells were lysed in RIPA buffer (Thermo Fisher) with protease inhibitor cocktail (Roche). Protein quantification was performed using a BCA Kit (Thermo Fisher). Protein lysate was subjected to SDS-PAGE and subsequently electrotransferred to a PVDF membrane (Bio-Rad) using wet transfer. Then primary antibodies and HRP conjugated secondary antibodies were incubated with the 5% milk blocked membrane. The imaging and quantification of target proteins was obtained by ChemiDoc MP imaging system (Bio-Rad).

### RT-qPCR

Total RNA was extracted using RNeasy Mini Plus Kit (Qiagen). Then One-step PrimeScript RT-PCR kit (Takara) was used to generate cDNA. RT-qPCR was performed with PowerUp SYBR Green Master Mix (Thermo Fisher) in QuantStudio 6 Pro Real-Time PCR System (Thermo Fisher). All primer sequences for qPCR are listed below:

SIRT6- Forward: 5′-ATGGAGCCTTCGGCTGACT-3′, Reverse: 5′-GTAACTATTCGGTGCGTTGGG-3′;

P16- Forward: 5′-ATGGAGCCTTCGGCTGACT-3′, Reverse: 5′- GTAACTATTCGGTGCGTTGGG-3′;

P21- Forward: 5′-CGATGGAACTTCGACTTTGTCA-3′, Reverse: 5′- GCACAAGGGTACAAGACAGTG-3′;

Lamin B1- Forward: 5′-GTAAGCACTGATTTCCATGTCCA-3′, Reverse: 5′- GAAAAAGACAACTCTCGTCGCA-3′;

IL6- Forward: 5′- ACTCACCTCTTCAGAACGAATTG-3′, Reverse: 5′- CCATCTTTGGAAGGTTCAGGTTG-3′;

### siRNA transfection

siRNAs targeting human USP10 (sc-365828) and Vimentin (sc-373717) were purchased from Santa Cruz Biotechnology. siRNAs were transfected into MSCs using Lipofectamine RNAiMAX Reagent (Thermo Fisher) according to the manufacturer’s instructions.

### Plasmids

For flag-VIM overexpression in MSC, VIM CDS were amplified from cDNA of MSC using primers (TGTCGTGACAAGTTTGTACAGCCACCATGGACTACAAAGACGATGACGACAAGTCCACCAGGTCCGTGTCCTCGTCCTCCTAC and TCGATTATCACCACTTTGTACATTATTCAAGGTCATCGTGATGCTGAGAAGT). Purified PCR products were cloned into pLV-Neo-EF1A backbone (VectorBuilder) using NEBuilder (NEB) according to the manufacturer’s instructions. Control plasmid expressing Flag tag was constructed by the insertion of annealed Flag sequences (GTACAGCCACCATGGACTACAAAGACGATGACGACAAGTAGT and GTACACTACTTGTCGTCATCGTCTTTGTAGTCCATGGTGGCT) into pLV-Neo-EF1A backbone using Quick Ligase (NEB).

For GFP-progerin overexpression in MSC, GFP-progerin CDS were amplified from pBABE-puro-GFP-progerin plasmids (Addgene, 17663) and cloned into pLenti4 backbone.

### Lentiviral preparation and cell transduction

To generate lentiviral particles, HEK293T cells were co-transfected with Plenti4-GFP-progerin, pLV-Neo-EF1A-Flag-VIM or pLV-Neo-EF1A-Flag, along with psPAX2 and pMD2.G (Addgene) packaging plasmids, using Lipofectamine 3000 (Thermo Fisher). Viral supernatants were collected 48 hours post-transfection, filtered through a 0.45 uM filter, and concentrated using the Lenti-X Concentrator (Takara) in accordance with the manufacturer’s protocol. Cells were transduced with the concentrated lentivirus in the presence of 10 µg/ml polybrene (Millipore). For GFP-progerin overexpression, cells were sorted using the BD Influx cell sorter after expansion one passages post-transduction to ensure all cells were successfully transduced. For VIM overexpression, cells were selected with 200 ng/ml G418 (Thermo Fisher) to establish stable neomycin-resistant populations for downstream analyses.

### Immunofluorescence staining

Cells were fixed in 4 % paraformaldehyde at room temperature (RT) for 15 min, permeabilized in 0.4% Triton X-100/PBS at RT for 10 min. After blocking with 10% donkey serum (Jackson ImmunoResearch Labs)/PBS for 1 h, cells were incubated with primary antibodies at 4 °C overnight and the corresponding secondary antibody (Invitrogen) at RT for 45 min. Nuclei were stained with Hoechst 33342 (Thermo Fisher 62249).

### RNA-seq library preparation

Total RNA of MSCs was isolated with the RNeasy Mini Plus Kit (Qiagen) following the manuals and then the integrity of RNA was checked by Bioanalyzer (Agilent). For hMSCs at early passages, RNA-seq libraries were constructed using NEBNext Ultra II RNA library Prep Kit and rRNA Depletion Kit (NEB) according to the manufacture’s protocol and paired-end reads were generated from Illumina NextSeq 550 at the Genomic Core of Albert Einstein College of Medicine. For hMSCs overexpressing GFP-progerin, RNA-seq libraries were constructed using Ribo-Zero Plus rRNA Depletion Kit (Illumina) according to the manufacture’s protocol and paired-end reads were generated from Illumina NovaSeq 6000 at Columbia University Genome Center.

### RNA-seq data processing

Trim Galore (0.6.10) was used to trim adaptor sequences and filter out low-quality reads. Processed reads were aligned to the human reference genome (hg38) using STAR (2.7.11b) and counted by featureCounts from Subread 2.0.2 package at gene level. Differential gene expression was analyzed by DESeq2 and assessed for Gene ontology or Gene Set Enrichment Analysis by clusterProfiler 4.0. TPM expression values were calculated using TPMCalculator.

### Transposable element expression analysis

To evaluate the expression levels of repetitive elements, the cleaned reads were mapped to the human reference genome (hg38) STAR software (version 2.7.11b) with the parameter --winAnchorMultimapNmax 100 --outFilterMultimapNmax 100 --outFilterMismatchNoverLmax 0.05. TE annotation file GRCh38_GENCODE_rmsk_TE.gtf was obtained from https://www.mghlab.org/software/tetranscripts. Transposable element quantification and the differential analysis were computed using the TEtranscripts software (version 2.2.3). In brief, TEtranscripts simultaneously counted the gene abundances and transposon abundances and utilized DESeq2 for the differential analysis.

### Statistical analysis

The statistical analyses were performed using PRISM software (Graphpad Software). Comparisons were performed with unpaired two-tail student’s t-test unless otherwise stated. P < 0.05 was defined as statistically significant.

## Data availability

All sequencing data were deposited to or fetched from Sequence Read Archive (SRA) or Gene Expression Omnibus (GEO) in the National Center for Biotechnology Information. The accession numbers were as follows: RNA-seq of SIRT6 hMSCs at early passage (PRJNA862192), RNA-seq of hMSCs expressing GFP-progerin (PRJNA1272066), RNA-seq of proliferating and senescent hMSCs (PRJNA1272555), RNA-seq of SIRT6-knockout hMSCs (GSE64642).

---

**Fig. S1.**
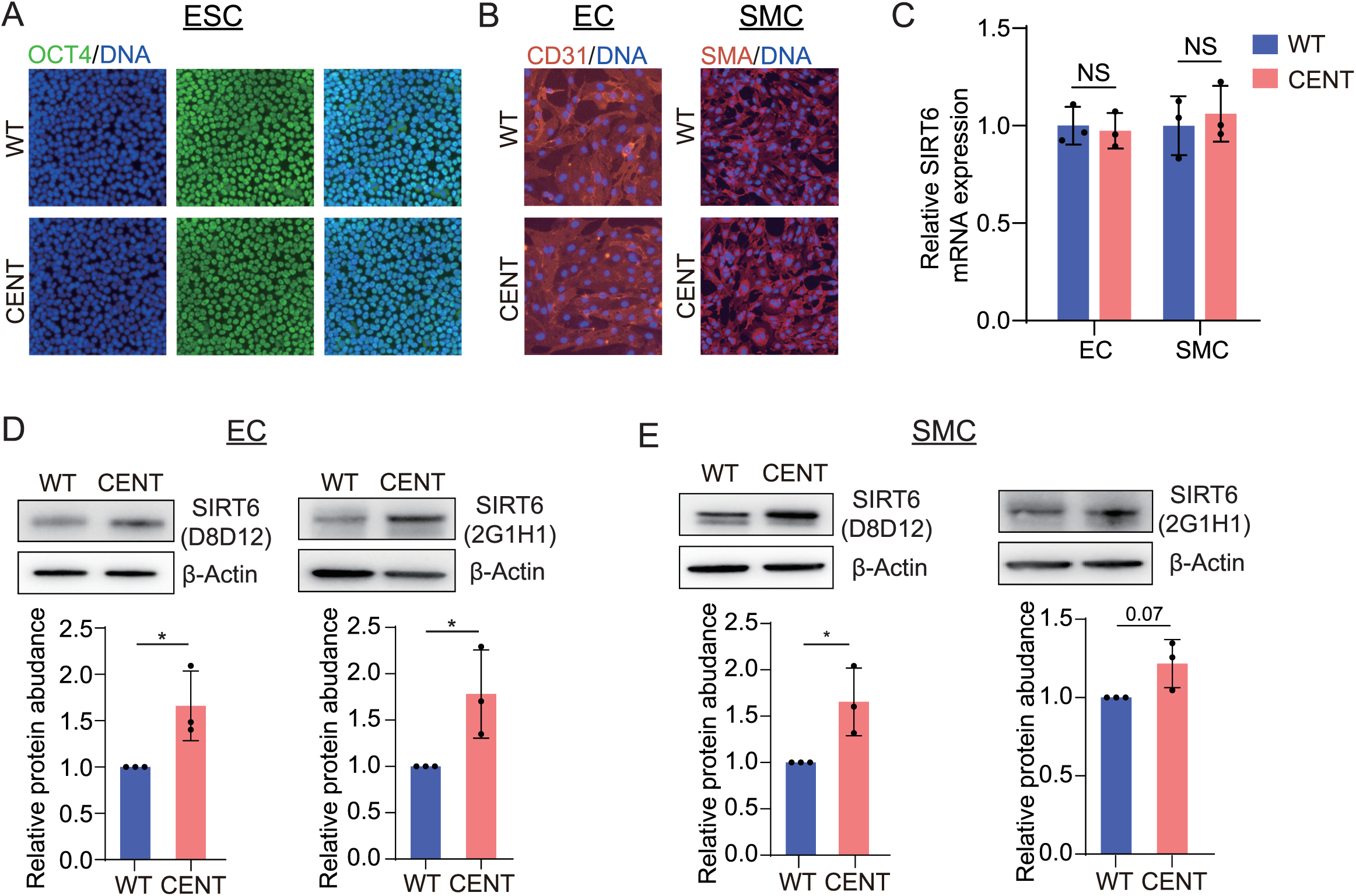
Elevated SIRT6 protein levels in hESCs and hESC-derived hECs and hSMCs carrying centenarian SIRT6 variants. **(A)** IF of pluripotency marker OCT4 in WT and CENT hESCs. **(B)** IF of EC marker CD31 and SMC marker SMA in hESC-derived somatic cells. **(C)** SIRT6 mRNA levels in WT and CENT hECs and hSMCs, gene expression was normalized to 18s rRNA; data were represented as mean ± SD; n = 3; NS, not significant. **(D)** SIRT6 protein levels in WT and CENT hECs. data were represented as mean ± SD; n = 3; *, p < 0.05. **(E)** SIRT6 protein levels in WT and CENT hSMCs. data were represented as mean ± SD; n = 3; *, p < 0.05.

**Fig. S2.**
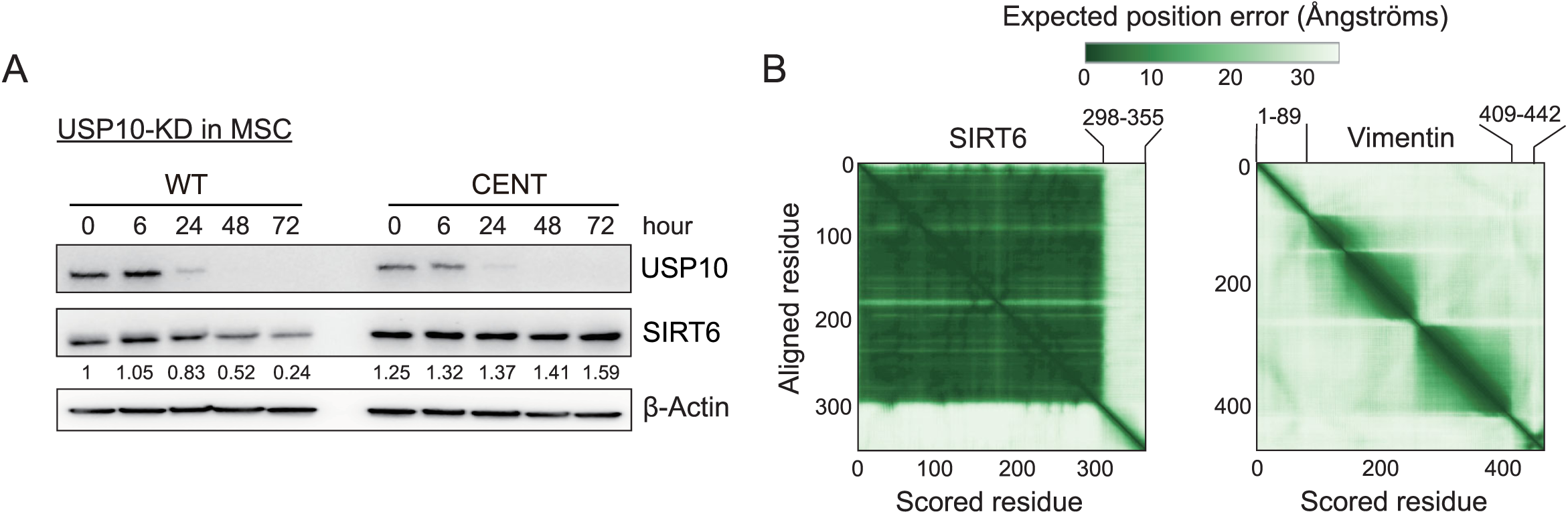
Vimentin-mediated elevation of SIRT6 protein levels in CENT hMSCs. **(A)** Protein levels of USP10 and SIRT6 in WT and CENT hMSCs transfected with siRNAs targeting USP10. **(B)** AlphaFold-predicted structural confidence for SIRT6 and VIM. Low confidence scores correspond to predicted intrinsic disorder regions (IDRs).

**Fig. S3.**
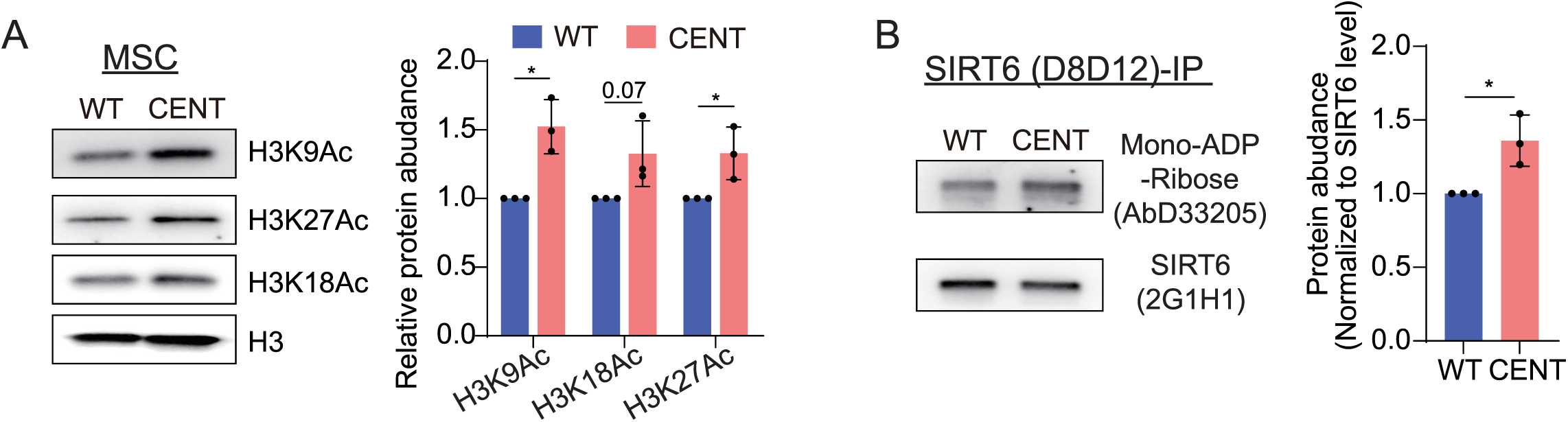
Centenarian SIRT6 variants alter enzymatic activities in a cellular context. **(A)** Protein levels of H3K9Ac, H3K27Ac and H3K18Ac in WT and CENT hMSCs. Protein levels were normalized to H3; data were represented as mean ± SD; n = 3; *, p < 0.05. **(B)** Relative proportion of mADP-ribosylated SIRT6 protein in hMSCs Endogenous SIRT6 was immunoprecipitated from WT and CENT hMSC lysates using anti-SIRT6 antibody (D8D12). IP samples were analyzed by western blotting with antibodies against SIRT6 (2G1H1) and mono-ADP-Ribose (AbD33205). Protein levels were normalized to SIRT6; data were represented as mean ± SD; n = 3; *, p < 0.05.

**Fig. S4.**
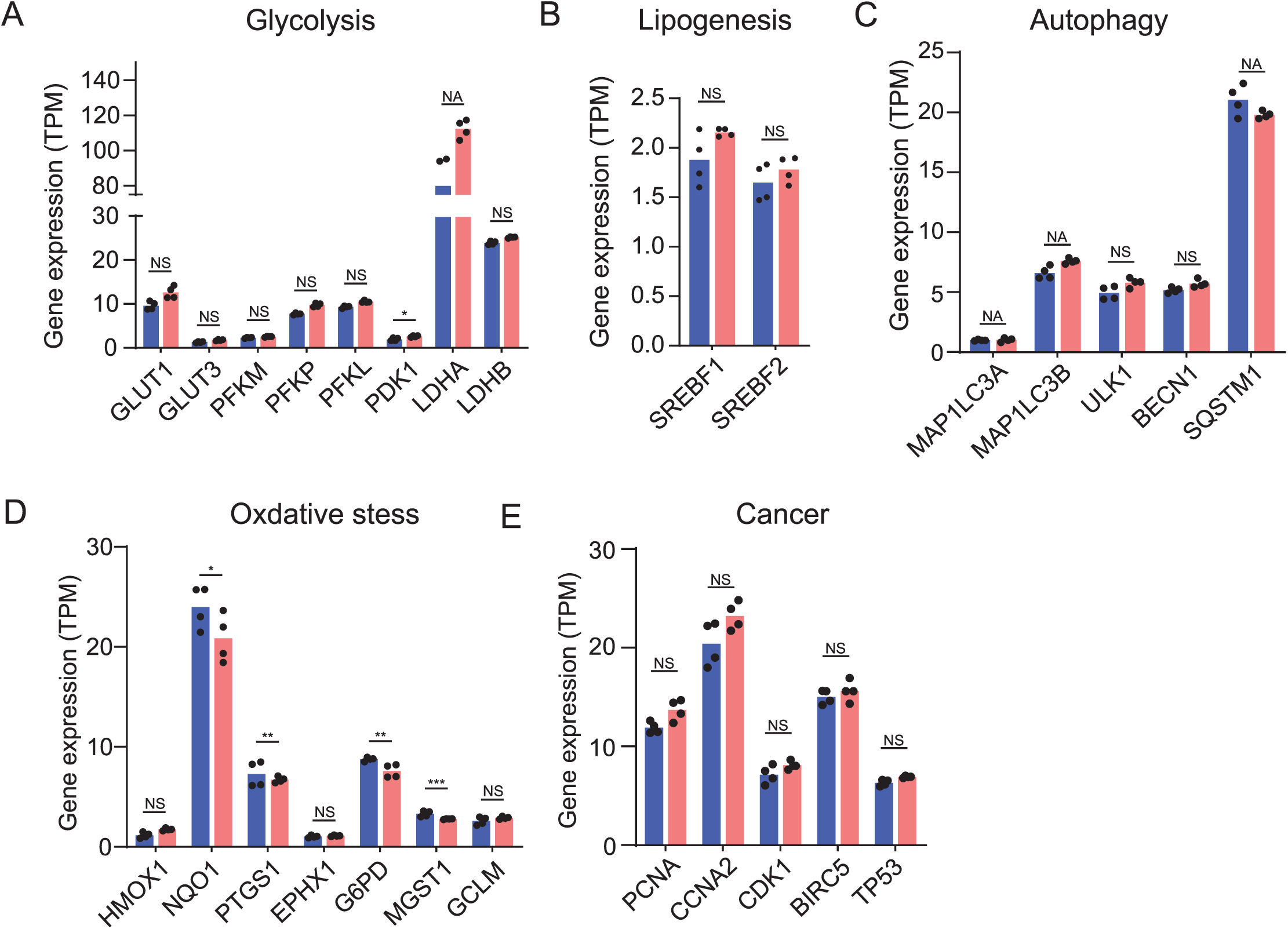
RNA-seq expression profiles of genes involving in pathways previously linked to SIRT6. **(A-E)** Pathways include glycolysis **(A)**, lipogenesis **(B)**, autophagy **(C)**, oxidative stress **(D)**, and cancer **(E)**. Normalized expression values (TPM: Transcripts Per Million) were used for plotting; data were represented as mean; Significance were indicated by adjusted. P value from DESeq2 analysis; *, p < 0.05; **, p < 0.01; ***, p < 0.001; NS, not significant; NA, not available.

**Fig. S5.**
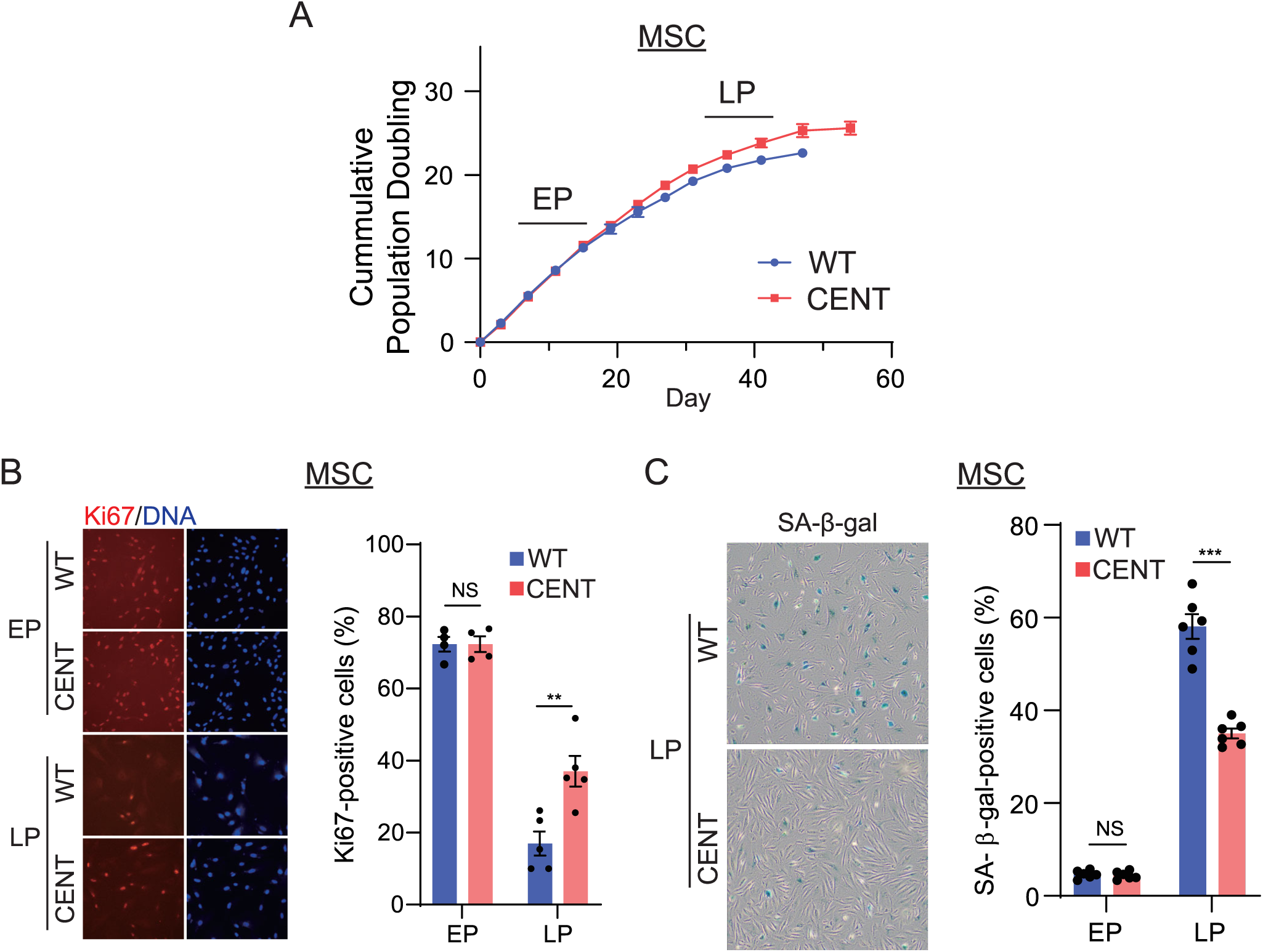
Centenarian SIRT6 variants delay replicative senescence. **(A)** Growth curves of WT and CENT hMSCs. EP (early passage, passage 3-5); LP (late passage, passage 10-11), data were represented as mean ± SD, n = 2. **(B)** IF staining of Ki67 in WT and CENT hMSCs at EP and LP. Quantification based on 5 independent images capturing over 500 nuclei; data were represented as mean ± SD; NS, not significant; **, p < 0.01. **(C)** SA-β-gal staining in WT and CENT hMSCs at EP and LP. Quantification based on 5 independent images capturing over 500 nuclei; data were represented as mean ± SD; NS, not significant; ***, p < 0.001.

## References

1 Kugel, S. & Mostoslavsky, R. Chromatin and beyond: the multitasking roles for SIRT6. Trends Biochem Sci 39, 72–81, doi:10.1016/j.tibs.2013.12.002 (2014).

2 Klein, M. A. & Denu, J. M. Biological and catalytic functions of sirtuin 6 as targets for small-molecule modulators. J Biol Chem 295, 11021–11041, doi:10.1074/jbc.REV120.011438 (2020).

3 Kanfi, Y. et al. The sirtuin SIRT6 regulates lifespan in male mice. Nature 483, 218–221, doi:10.1038/nature10815 (2012).

4 Roichman, A. et al. Restoration of energy homeostasis by SIRT6 extends healthy lifespan. Nat Commun 12, 3208, doi:10.1038/s41467-021-23545-7 (2021).

5 Tian, X. et al. SIRT6 Is Responsible for More Efficient DNA Double-Strand Break Repair in Long-Lived Species. Cell 177, 622–638 e622, doi:10.1016/j.cell.2019.03.043 (2019).

6 Simon, M. et al. A rare human centenarian variant of SIRT6 enhances genome stability and interaction with Lamin A. EMBO J 41, e110393, doi:10.15252/embj.2021110393 (2022).

7 Fehrer, C. & Lepperdinger, G. Mesenchymal stem cell aging. Exp Gerontol 40, 926–930, doi:10.1016/j.exger.2005.07.006 (2005).

8 Al-Azab, M., Safi, M., Idiiatullina, E., Al-Shaebi, F. & Zaky, M. Y. Aging of mesenchymal stem cell: machinery, markers, and strategies of fighting. Cell Mol Biol Lett 27, 69, doi:10.1186/s11658-022-00366-0 (2022).

9 Zhang, W. et al. Aging stem cells. A Werner syndrome stem cell model unveils heterochromatin alterations as a driver of human aging. Science 348, 1160–1163, doi:10.1126/science.aaa1356 (2015).

10 Jing, Y. et al. Genome-wide CRISPR activation screening in senescent cells reveals SOX5 as a driver and therapeutic target of rejuvenation. Cell Stem Cell 30, 1452–1471 e1410, doi:10.1016/j.stem.2023.09.007 (2023).

11 Wang, W. et al. A genome-wide CRISPR-based screen identifies KAT7 as a driver of cellular senescence. Sci Transl Med 13, doi:10.1126/scitranslmed.abd2655 (2021).

12 Thirumurthi, U. et al. MDM2-mediated degradation of SIRT6 phosphorylated by AKT1 promotes tumorigenesis and trastuzumab resistance in breast cancer. Sci Signal 7, ra71, doi:10.1126/scisignal.2005076 (2014).

13 Ronnebaum, S. M., Wu, Y., McDonough, H. & Patterson, C. The ubiquitin ligase CHIP prevents SirT6 degradation through noncanonical ubiquitination. Mol Cell Biol 33, 4461–4472, doi:10.1128/MCB.00480-13 (2013).

14 Lin, Z. et al. USP10 antagonizes c-Myc transcriptional activation through SIRT6 stabilization to suppress tumor formation. Cell Rep 5, 1639–1649, doi:10.1016/j.celrep.2013.11.029 (2013).

15 Basu, A. et al. Vimentin undergoes liquid-liquid phase separation to form droplets which wet and stabilize actin fibers. Proc Natl Acad Sci U S A 122, e2418624122, doi:10.1073/pnas.2418624122 (2025).

16 Holehouse, A. S. & Kragelund, B. B. The molecular basis for cellular function of intrinsically disordered protein regions. Nat Rev Mol Cell Biol 25, 187–211, doi:10.1038/s41580-023-00673-0 (2024).

17 Jumper, J. et al. Highly accurate protein structure prediction with AlphaFold. Nature 596, 583–589, doi:10.1038/s41586-021-03819-2 (2021).

18 Ginell, G. M. et al. Sequence-based prediction of intermolecular interactions driven by disordered regions. Science 388, eadq8381, doi:10.1126/science.adq8381 (2025).

19 Pan, H. et al. SIRT6 safeguards human mesenchymal stem cells from oxidative stress by coactivating NRF2. Cell Res 26, 190–205, doi:10.1038/cr.2016.4 (2016).

20 Zhang, Z. et al. Increased hyaluronan by naked mole-rat Has2 improves healthspan in mice. Nature 621, 196–205, doi:10.1038/s41586-023-06463-0 (2023).

21 Sebastian, C. et al. The histone deacetylase SIRT6 is a tumor suppressor that controls cancer metabolism. Cell 151, 1185–1199, doi:10.1016/j.cell.2012.10.047 (2012).

22 Kim, H. S. et al. Hepatic-specific disruption of SIRT6 in mice results in fatty liver formation due to enhanced glycolysis and triglyceride synthesis. Cell Metab 12, 224–236, doi:10.1016/j.cmet.2010.06.009 (2010).

23 Iachettini, S. et al. Pharmacological activation of SIRT6 triggers lethal autophagy in human cancer cells. Cell Death Dis 9, 996, doi:10.1038/s41419-018-1065-0 (2018).

24 Shao, J. et al. Autophagy induction by SIRT6 is involved in oxidative stress-induced neuronal damage. Protein Cell 7, 281–290, doi:10.1007/s13238-016-0257-6 (2016).

25 Kawahara, T. L. et al. Dynamic chromatin localization of Sirt6 shapes stress- and aging-related transcriptional networks. PLoS Genet 7, e1002153, doi:10.1371/journal.pgen.1002153 (2011).

26 Kubben, N. & Misteli, T. Shared molecular and cellular mechanisms of premature ageing and ageing-associated diseases. Nat Rev Mol Cell Biol 18, 595–609, doi:10.1038/nrm.2017.68 (2017).

27 Kubben, N. et al. Repression of the Antioxidant NRF2 Pathway in Premature Aging. Cell 165, 1361–1374, doi:10.1016/j.cell.2016.05.017 (2016).

28 Chojnowski, A. et al. Heterochromatin loss as a determinant of progerin-induced DNA damage in Hutchinson-Gilford Progeria. Aging Cell 19, e13108, doi:10.1111/acel.13108 (2020).

29 Della Valle, F. et al. LINE-1 RNA causes heterochromatin erosion and is a target for amelioration of senescent phenotypes in progeroid syndromes. Sci Transl Med 14, eabl6057, doi:10.1126/scitranslmed.abl6057 (2022).

30 Liu, X. et al. Resurrection of endogenous retroviruses during aging reinforces senescence. Cell 186, 287–304 e226, doi:10.1016/j.cell.2022.12.017 (2023).

31 Van Meter, M. et al. SIRT6 represses LINE1 retrotransposons by ribosylating KAP1 but this repression fails with stress and age. Nat Commun 5, 5011, doi:10.1038/ncomms6011 (2014).

32 Simon, M. et al. LINE1 Derepression in Aged Wild-Type and SIRT6-Deficient Mice Drives Inflammation. Cell Metab 29, 871–885 e875, doi:10.1016/j.cmet.2019.02.014 (2019).

33 Robbins, P. et al. Fucoidans are senotherapeutics that enhance SIRT6-dependent DNA repair. Res Sq, doi:10.21203/rs.3.rs-6613032/v1 (2025).

34 Biashad, S. A. et al. SIRT6 activator fucoidan extends healthspan and lifespan in aged wild-type mice. bioRxiv, 2025.2003.2024.645072, doi:10.1101/2025.03.24.645072 (2025).

35 Lin, J. R. et al. Rare genetic coding variants associated with human longevity and protection against age-related diseases. Nat Aging 1, 783–794, doi:10.1038/s43587-021-00108-5 (2021).

36 Gallagher, M. D. & Chen-Plotkin, A. S. The Post-GWAS Era: From Association to Function. Am J Hum Genet 102, 717–730, doi:10.1016/j.ajhg.2018.04.002 (2018).

37 Geng, L. et al. Chemical screen identifies a geroprotective role of quercetin in premature aging. Protein Cell 10, 417–435, doi:10.1007/s13238-018-0567-y (2019).

38 Biskou, O. et al. The type III intermediate filament vimentin regulates organelle distribution and modulates autophagy. PLoS One 14, e0209665, doi:10.1371/journal.pone.0209665 (2019).

39 Hirata, Y. et al. Vimentin binds IRAP and is involved in GLUT4 vesicle trafficking. Biochem Biophys Res Commun 405, 96–101, doi:10.1016/j.bbrc.2010.12.134 (2011).

40 Jiu, Y. Vimentin intermediate filaments function as a physical barrier during intracellular trafficking of caveolin-1. Biochem Biophys Res Commun 507, 161–167, doi:10.1016/j.bbrc.2018.10.199 (2018).

41 Lei, J. et al. Senescence-resistant human mesenchymal progenitor cells counter aging in primates. Cell 188, 5039–5061 e5035, doi:10.1016/j.cell.2025.05.021 (2025).

42 Yang, J. et al. Genetic enhancement in cultured human adult stem cells conferred by a single nucleotide recoding. Cell Res 27, 1178–1181, doi:10.1038/cr.2017.86 (2017).

43 Patsch, C. et al. Generation of vascular endothelial and smooth muscle cells from human pluripotent stem cells. Nat Cell Biol 17, 994–1003, doi:10.1038/ncb3205 (2015).

